# A stress-reduced passaging technique improves the viability of human pluripotent cells

**DOI:** 10.1101/2021.10.12.464142

**Authors:** Kazutoshi Takahashi, Chikako Okubo, Michiko Nakamura, Mio Iwasaki, Yuka Kawahara, Tsuyoshi Tabata, Yousuke Miyamoto, Knut Woltjen, Shinya Yamanaka

## Abstract

Xeno-free culture systems have expanded the clinical and industrial application of human pluripotent stem cells (PSCs). However, yet some problems, such as the reproducibility among the experiments, remain. Here we describe an improved method for the subculture of human PSCs. The revised method significantly enhanced the viability of human PSCs by lowering DNA damage and apoptosis, resulting in more efficient and reproducible downstream applications such as gene editing, gene delivery, and directed differentiation. Furthermore, the method did not alter PSC characteristics after long-term culture and attenuated the growth advantage of abnormal subpopulations. This robust passaging method minimizes experimental error and reduces the rate of PSCs failing quality control of human PSC research and application.

**Highlights:**

- The revised passaging method significantly increases the viability of human PSCs.
- The method triggers less DNA damage and apoptosis signals compared to the conventional method.
- The stress-reduced method improves the results of downstream applications.
- The method does not alter PSC characters and attenuates the overgrowth of abnormal subpopulations.

## Introduction

Xeno-free culture methods for human pluripotent stem cells (PSCs) are widespread. They allow the maintenance of human PSCs, such as embryonic stem cells (ESCs) and induced PSCs (iPSCs), without losing their differentiation potential. The development and optimization of culture vessel-coating reagents, such as synthetic substrates (Melkoumian et al., 2010; Villa-Diaz et al., 2010) and extracellular matrices (Rodin et al., 2010), have contributed to efficient culture systems without the inclusion of animal-derived components, including feeder cells and the Engelbreth-Holm-Swarm mouse tumor-derived basement membrane matrix (i.e., Matrigel) (Thomson et al., 1998; Xu et al., 2001). Particularly, recombinant extracellular matrices, such as laminin and vitronectin, allow the rapid and efficient adhesion of undifferentiated PSCs, which express compatible integrin receptors (Chen et al., 2011; Miyazaki et al., 2012; Nakagawa et al., 2014; Rodin et al., 2010; Xu et al., 2001).

Combining such fine-tuned culture systems with inhibitors of cell death, such as the Rho-associated protein kinase (ROCK) inhibitor, dramatically improves the survival of human PSCs during passage and makes single-cell culture of human PSCs possible (Chen et al., 2021; Watanabe et al., 2007). Single-cell cloning following gene disruption or correction by CRISPR/Cas9 facilitates the isolation of PSCs for in vitro disease modeling and as possible sources for cell therapies (Hotta and Yamanaka, 2015). In this way, single-cell culture of human PSCs is a promising method for the systematic quality control of clinical-grade cell lines and reproducibility of basic research.

On the other hand, unlike advances in the extracellular environment, the passaging procedure has not changed significantly in the last decade. Conventional passaging methods for human PSCs have adhered fundamentally to the classical procedure of dissociating cells after removing/neutralizing the enzymatic activity and/or a chelating reaction by adding growth media (Beers et al., 2012; Chen et al., 2011; Miyazaki et al., 2012; Nakagawa et al., 2014). Considering that the PSC culture conditions have significantly changed, there is room for optimizing the passaging procedure to fit current culture protocols. In this study, we revised a passage protocol for human PSCs cultured in xeno-free conditions. The method significantly improves the reproducibility of experiments and cell viability. Such controllable passaging facilitated the results of several downstream assays, such as gene editing, gene delivery, and differentiation. The modification of the passaging procedure did not change the PSC characteristics but did attenuate the growth advantage of abnormal subpopulations. The revised method promises to make PSC cultures safer and further expands the range of human PSC usage.

## Results

The conventional method of passaging human PSCs on a xeno-free culture described previously (Fig. 1A) (Nakagawa et al., 2014) results in poor reproducibility such as wide range of cell viability (Fig. 1B). To resolve this issue, we modified the procedure of cell passaging and explored the effects on cell viability in detail using two independent iPSC lines, WTB6 (Miyaoka et al., 2014) and WTC11 (Kreitzer et al., 2013), maintained on laminin-511 E8 fragment in a chemically defined media, StemFit (Miyazaki et al., 2012; Nakagawa et al., 2014) (Fig. 1A). First, we used TrypLE, 5 mM EDTA, or AccuMax (Kim et al., 2016) as the detachment reagent instead of a 1:1 mix of TrypLE and 0.5 mM EDTA (hereafter T1E1), which is used in the conventional method (Miyazaki et al., 2017; Nakagawa et al., 2014). Second, the incubation time was extended from 5 min to 10 min. Finally, the dissociation to single cells was performed directly in the detachment solution rather than replacing the detachment solution with growth media and dislodging the cells from the culture vessel using a cell scraper prior to dissociation. Using this revised method, we could easily detach the cells with all three detachment reagents by gently pipetting up and down. This was nearly impossible with the conventional protocol, which required the use of a cell scraper. As a result of the modifications, the revised method significantly and reproducibly improved the viability of human PSCs (Figs. 1B, S1A). As well as WTB6 and WTC11, the method improved the viability of the iPSC lines such as 201B7 (Takahashi et al., 2007) and 585A1 (Okita et al., 2011), and the ESC line H9 (Thomson et al., 1998) more than 95% on average (Fig. S1B). The revised method also worked well for another xeno-free PSC culture condition combining recombinant vitronectin and Essential 8 media (Chen et al., 2011) (Fig. S1C). Since EDTA has been employed for the detachment of human PSCs from laminin-511 or vitronectin-coated culture vessels, we tested 5 mM EDTA instead of TrypLE in the revised method (Beers et al., 2012; Chen et al., 2011; Miyazaki et al., 2012). After 10 minutes of incubation and then dissociation in 5 mM EDTA, the cells were easily detached using a micropipette and showed high viability (Fig. 1B). In 0.5-5 mM EDTA, no apparent cytotoxicity was detected (Fig. S1D), although TrypLE but not EDTA generated single-cell suspensions efficiently by pipetting (Fig. S1E). This observation suggests that we can choose TrypLE or EDTA according to the type of downstream assay. Taken together, these data indicate higher viability of human PSCs in the passaging process.

**Fig. 1.**
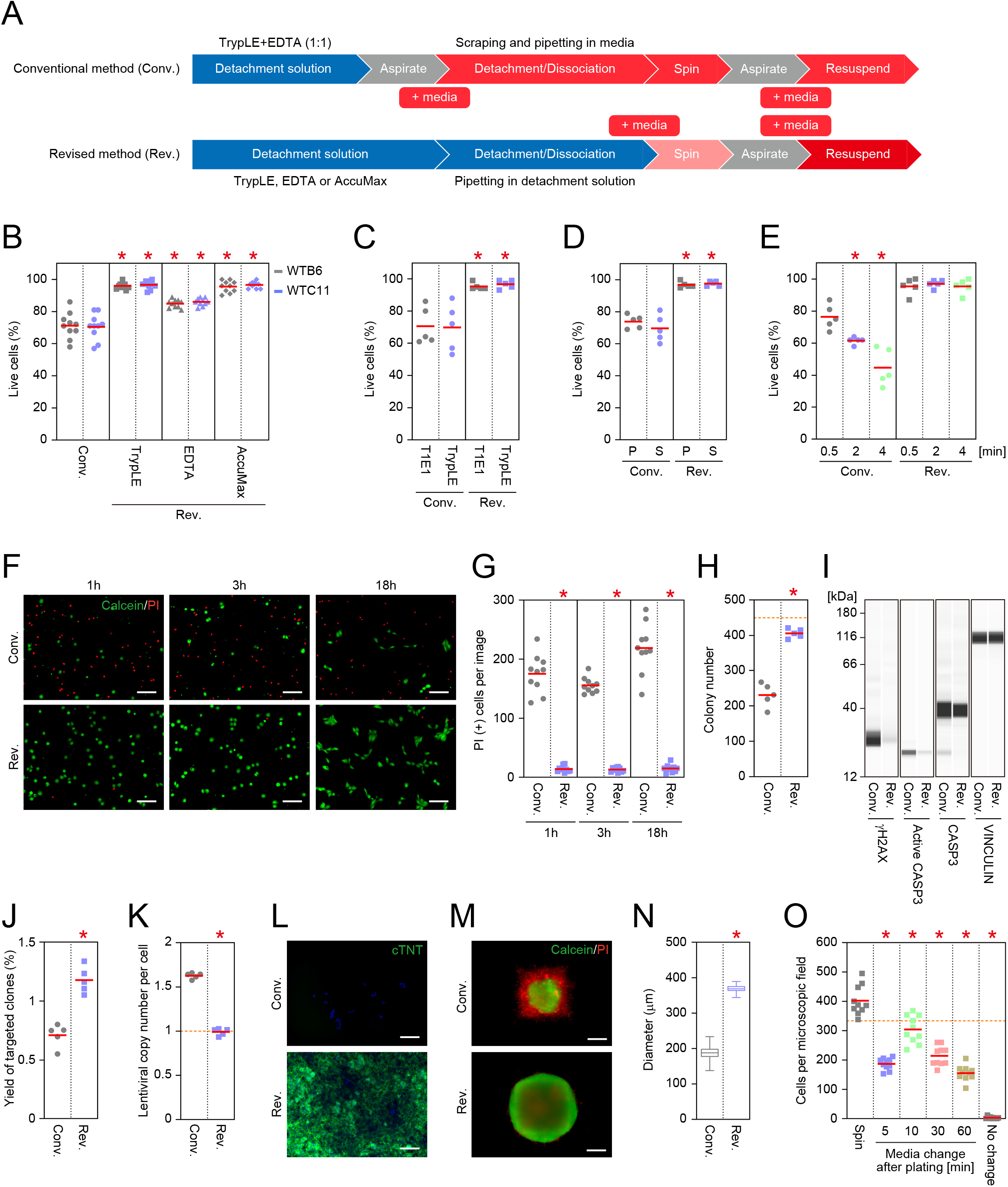
The revised passaging method improves the viability of human PSCs. A. A flow diagram of the cell passage methods used in the study. B. The viability of human PSCs after harvesting using the conventional (Conv.) or revised passaging method. *p<0.05 vs. Conv., one-way ANOVA (Tukey’s multiple comparisons test). n=10. C. The viability of human PSCs after harvesting using the conventional or revised passaging method with T1E1 or TrypLE. *p<0.05 vs. Conv., one-way ANOVA (Tukey’s multiple comparisons test). n=5. D. The viability of human PSCs after dissociation in growth media (Conv.) or dissociation reagent (Rev.) by pipetting (P) or using a cell scraper (S). *p<0.05 vs. Conv., one-way ANOVA (Tukey’s multiple comparisons test). n=5. E. The viability of human PSCs harvested after incubation in dissociation reagent or growth media for the indicated times. *p<0.05 vs. 0.5 min, one-way ANOVA (Tukey’s multiple comparisons test). n=5. F. Cell death after plating. Live and dead cells were visualized with Calcein-AM (green) and propidium iodide (PI, red), respectively. Bars, 100 μm. G. Quantification of dead cells after the indicated times of plating. Ten microscopic views were randomly imaged in each condition and time point, and the number of PI-stained dead cells were counted. *p<0.05 vs. Conv., one-way ANOVA (Tukey’s multiple comparisons test). H. The number of colonies derived from PSCs harvested using the conventional or revised method. The mandarin-colored broken line indicates 100% adhesion efficiency. *p<0.05 vs. Conv., unpaired t-test. n=5. I. Western blotting of DNA damage and cell death markers. Shown are the virtual blots of phospho-H2AX (γH2AX), cleaved Caspase-3 (Active CASP3), Caspase-3 (CASP3) and Vinculin in cells harvested using the conventional or revised method. J. The yield of AAVS1-targeted colonies. *p<0.05 vs. Conv., unpaired t-test. n=5. K. The copy number of lentiviral integrations in the genome per cell. *p<0.05 vs. Conv., unpaired t-test. n=5. L. Cardiac differentiation. Representative images of cardiac Troponin-T staining (cTNT, green) are shown. Nuclei were visualized with Hoechst 33342 (blue). Bars, 100 μm. M. Neurosphere formation. Representative images of Calcein/PI-stained neurospheres after 18 h of plating in AggreWell800. Bars, 100 μm. N. The diameters of spheres labeled with Calcein derived from PSCs harvested using the conventional (n=116) or revised (n=114) method were measured. The boxes and whiskers indicate the mean and the minimum to maximum values, respectively. *p<0.05 vs. Conv., unpaired t-test. O. The effects of TrypLE on the adhesion of PSCs. Cells harvested with the revised method were plated in media containing TrypLE. The media was changed at the indicated times after plating or not at all (no change). Microscopic views under a 10x objective were randomly chosen, and the number of nuclei labeled by Hoechst 33342 were counted. *p<0.05 vs. Spin., one-way ANOVA (Tukey’s multiple comparisons test). n=10.

Next, to elucidate which part of the passaging is crucial for viability, we systematically tested the detachment and dissociation processes. First, we detached human PSCs using TrypLE or T1E1 and then harvested the cells in detachment solution or growth media. We found that the cell viability depended on the condition of the cell dissociation process regardless of the components of the detachment solution (Fig. 1C). Using a cell scraper in detachment solution did not decrease the viability, suggesting that damage induced by scraping is not responsible for the cell death in the conventional method (Fig. 1D). The data shown in Figs. 1C and 1D led us to hypothesize that the enforced cell tearing after replacing the detachment reagent with growth media decreased the cell viability. Indeed, after replacing the detaching solution with growth media, the cells immediately began to re-attach to the culture vessels, leading to difficult and inefficient cell detachment by pipetting (Fig. S1F). Moreover, a longer incubation time in growth media before cell scraping significantly decreased cell viability (Fig. 1E). On the other hand, prolonged incubation of human PSCs in TrypLE solution did not affect the viability (Figs. 1E, S1G). All in all, we concluded that cell dissociation in growth media is a critical step for cell viability.

We found that the revised method not only improves the cell viability immediately after harvesting but also significantly improves human PSCs adhesion to the culture vessel upon plating compared to cells collected using the conventional method if the same number of live cells are plated (Figs 1F, 1G, S1H, S1I). Plating the cells at higher density (1 × 10^5^ cells/cm^2^) did not change this tendency (Fig. S1J). To quantify the cell adhesion efficiency by eliminating the influence of cell division, we plated the cells at clonal density and counted the number of colonies. As a result, the cells harvested by the revised method formed colonies with 90.2±2.85% adhesion efficiency whereas those harvested by the conventional method had 51.2±7.34% efficiency, which is similar to a previous report (Fig. 1H) (Miyazaki et al., 2012). Given that plating the same number of live cells just after harvesting by different methods made such a significant difference in the adhesion efficiency, these data suggest that further cell death after plating is a cause of compromised plating efficiency. A western blot analysis revealed that a DNA damage marker, phosphorylated Ser139 of histone H2AX variant (γH2AX), and an apoptosis marker, cleaved Caspase-3, were significantly increased in cells harvested by the conventional method compared to the revised method (Fig. 1I). These data suggest that the revised passaging method improves the viability and adhesion efficiency of human PSCs by not inducing DNA damage or apoptosis.

Based on these results, we thought that such a robust passage could improve the efficiency and accuracy of downstream experiments. First, we tested if the revised passaging method increases the yield of clones gene-edited by a CRISPR/Cas9 platform. As expected, the cells harvested using the revised method prior to electroporation generated higher efficiency when targeting the Adeno-associated virus integration site 1 (AAVS1) locus than the conventional method (Fig. 1J). Next, we examined if we could control the lentiviral infection efficiency that is often required for the transduction of small guide RNA library in CRISPR-based high-throughput screening (Gilbert et al., 2014; Horlbeck et al., 2016; Koike-Yusa et al., 2014). Cells harvested using either method were exposed to the lentivirus at a multiplicity of infection = 1. As a result, the cells yielded by the conventional method exhibited higher copies (1.63±0.03 per cell) of lentiviral integration due to lower cell numbers for the infection. On the other hand, the cells harvested by the revised method were nearly single copy, as expected (0.99±0.04 per cell) (Fig. 1K). Furthermore, we hypothesized that plating an accurate number of cells with high viability facilitates the differentiation of PSCs. Again, as expected, the revised method yielded more Troponin T (+) cardiomyocytes by direct differentiation than cells harvested by the conventional method, which instead caused massive cell death or poor differentiation (Fig. 1L) (Lian et al., 2012). Moreover, the PSCs harvested by the revised method efficiently formed neurospheres with few dead cells (Figs. 1M, S1K) and could produce larger and more uniform sized spheres than the cells harvested by the conventional method (Fig. 1N). These data suggest that the revised passaging method makes downstream applications using human PSCs more efficient and controllable.

Because the revised method dissociates and collects the cells in the detachment reagent, we spun the cells down to remove the supernatant and resuspended the cell pellet in growth media (Fig. 1A). The cells suspended in media containing 10% TrypLE (volume) floated after 18 h of plating, suggesting that removing the detachment reagent is needed (Fig. 1O). Considering our intentions to apply the method to automated and/or high-throughput cell culture systems for industrial applications and clinical use, excluding the centrifuge process would be more practical. Therefore, taking advantage of the characteristic that human PSCs rapidly adhere to the culture vessel in the presence of a suitable extracellular matrix, such as laminin-511, we tested if media containing the 10% of detachment reagent can be replaced with complete growth media after plating. Indeed, the number of attached cells after 18 h of plating decreased one half by changing the media 5 min after plating (Fig. 1O). Noteworthy, changing the media 10 min after plating resulted in a 76.5±14.0% yield based on the number of cells suspended in complete media before plating. The lost quarter of adhered cells is significant, but the result is still much better than the number of adhered cells harvested by the conventional method before plating (Figs. 1O, S1I). Taken together, the revised passaging method is potentially beneficial for the preparation of human PSCs to be used in later applications.

One of the highest priority problems for PSC culture systems is avoiding malignant transformation by the abnormal growth of unexpected subpopulations. Thus, we created a human iPSC line overexpressing BCL2L1, which drives the strong selective advantage of chromosome 20q11.21 amplification (Amps et al., 2011; Avery et al., 2013), and tested the effect of the passage method on its growth rate. The revised passaging method significantly attenuated the growth advantage of BCL2L1-expressing iPSCs, whereas the conventional method biased amplification (Figs. 2A, 2B). This observation suggests that the revised method keeps human PSCs stable during continuous culture. We validated human PSCs split 30 times over 100 days using the revised method. These cells kept expressing core PSC transcription factors, such as OCT3/4 (Okamoto et al., 1990; Scholer et al., 1990) and NANOG (Chambers et al., 2003; Mitsui et al., 2003), and displayed an affinity for rBC2LCN (Onuma et al., 2013) (Figs. 2C, S2A). We confirmed that passaging with the revised method did not induce major changes of global RNA expression, the expression of PSC and differentiation-deficiency (DD) markers (Koyanagi-Aoi et al., 2013; Ohnuki et al., 2014) and global protein expression levels (Figs. 2D, 2E, S2B, S2C) and that the cells could differentiate into three germ layers through directed differentiation methods (Figs. 2F, 2G, S2D, S2E) and have no apparent karyotype abnormalities (Figs. 2H, S2F). Altogether, these data suggest that the revised passaging method is suitable for human PSC subculture.

**Fig. 2.**
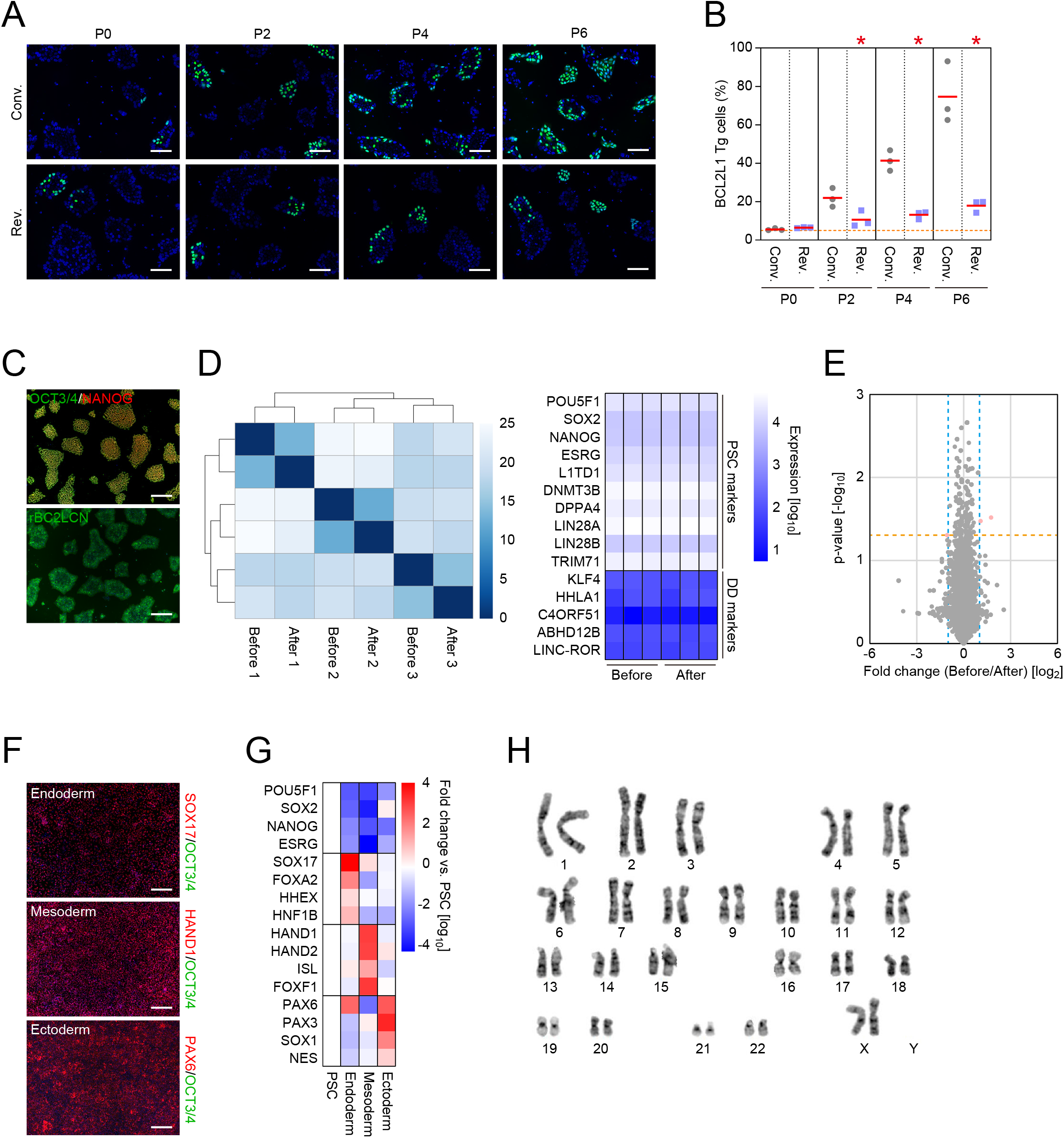
The revised passaging method does not alter PSC characteristics. A. Representative images of the competition between wild-type (Clover negative) and BCL2L1 transgenic (Clover positive) cells at the indicated passage numbers. Nuclei were visualized with Hoechst 33342 (blue). Bars, 100 μm. B. The percentage of BCL2L1 transgenic (Tg) cells during the competition culture measured by genomic PCR. The mandarin-colored broken line indicates the percentage of BCL2L1 Tg cells before the competition. *p<0.05 vs. Conv., one-way ANOVA (Tukey’s multiple comparisons test). n=3. C. PSC markers in WTB6 iPSCs after 30 passages with the revised method. Nuclei were visualized with Hoechst 33342 (blue). Bars, 200 μm. D. The RNA expression. Shown are clustering analysis of global RNA expression (left) and the PSC and differentiation-deficiency (DD) markers between WTB6 iPSCs before and after ≥30 passages with the revised method. n=3. E. The protein expression. The volcano plot shows the comparison of global protein expression between WTB6 iPSCs before and after cultivation with passaging using the revised method (>30 passages). n=3. The mandarin-colored and turquoise-colored broken lines indicate p=0.05 and fold change=2, respectively. Red dots indicate differentially expressed proteins such as CNEP1R1, MATK and Trypsin. F. Directed differentiation into trilineage. Shown are representative immunocytochemistry images of differentiated cells stained for each lineage marker, including SOX17, HAND1 and PAX6 (red), and a PSC marker, OCT3/4 (green). Nuclei were visualized with Hoechst 33342 (blue). Bars, 200 μm. G. The expression of lineage markers. The heatmap shows the relative expression of PSC and lineage markers in directed differentiated cells compared to parental iPSCs. n=3. H. A representative G-banding image shows normal 46XX karyotype.

## Discussion

In this study, we provide a revised passaging method optimized for the xeno-free culture of human PSCs. The major change compared with the conventional method was cell dissociation prior to media replacement; other factors, such as reagents and mechanical dissociation techniques, are flexible. Since human PSCs rapidly adhere to the matrix-coated culture vessel again after replacing dissociation reagent with growth media, forced cell detachment by scraping or pipetting should be stressful (Figs. 1I, S1F). However, we demonstrated that quick cell dissociation after adding growth media achieves better viability (Fig. 1E). Although this quick dissociation may be applicable for one-by-one treatment of cultured cells, multi-throughput experiments need a more flexible time window of the dissociation step. Using the revised method, we showed that 10 min of incubation in TrypLE at 37°C for detachment and a subsequent 60 min of incubation at room temperature did not affect the viability of human PSCs (Fig. S1G). Therefore, we concluded that the revised method is suitable for the multi- or high-throughput treatment of human PSC culture.

On the other hand, the continuous commingling of TrypLE with the growth media overnight detached human PSCs from the culture vessel, suggesting that TrypLE must be removed. The ideal solution for cell viability then is replacing the TrypLE-containing media with complete media after spinning the cells down. For an automated cell culture system, however, the centrifuging process is preferred to be excluded. We showed a model of the spin-free method by changing the media after plating the cells onto a culture vessel. Removing the TrypLE-containing media after 10 minutes of plating yielded 75% of cells after 18 h of plating compared to spinning the cells down before plating. While there may still be a margin for improvement, this approach is nevertheless an attractive model for PSC-based applications.

The use of ROCK inhibitor in the human PSC culture increases the cloning efficiency from less than 1% to 25% (Watanabe et al., 2007). Additionally, the high affinity between integrin substrates and human PSCs allows for a viability of around 50% (Beers et al., 2012; Chen et al., 2011; Miyazaki et al., 2012). The stress-reduced passaging method shown in this study enables 90% adhesion efficiency even at clonal density. This study optimized the passaging procedure, which has been relatively ignored compared with the media and matrix in efforts to optimize xeno-free PSC cultures. That improving the passaging method was sufficient to reach a yield with more than 95% viability suggests the proper combination of available xeno-free media and recombinant matrix is already effective for efficient culturing. Since our revised method simply modifies the cell dissociation condition, it can be applied to current procedures without significant modifications and will likely improve the results of basic PSC research and applications.

### Limitations of the study

The protocol provided in this study is suitable for PSCs maintained on recombinant extracellular matrices that allow for rapid and efficient adhesion. This feature overcomes the issue of forcibly dislodging PSCs after adding growth media, which reduces their viability. Since classical culture methods that use Matrigel or feeder cells do not have this dislodgement issue, we believe that our revised protocol has few advantages. The revised method enhances the viability of cells only during passaging and not during growth. Adding cell death inhibitors is still needed to improve the expansion efficiency (Chen et al., 2021; Watanabe et al., 2007).

Finally, dissociation with EDTA led to a lower viability than with TrypLE or AccuMax (Figs. 1B, S1C, S1D). However, as shown in Fig. S1E, EDTA treatment did not produce single cells efficiently. Thus, the cell viability can be underestimated because of selective single cell counting. Therefore, we cannot conclude an advantage of enzymatic treatment compared to EDTA.

## Acknowledgments

We thank B. Conklin, K. Okita, D. Trono and J. Weissman for sharing materials, K. Higashi, K. Kamegawa, R. Kato, M. Saito, and S. Takeshima for administrative support, and P. Karagiannis and T. Matsumoto for reading the manuscript. This work was supported by Grants-in-Aid for Scientific Research (20K20585 and 21H02155) from the Japanese Society for the Promotion of Science (JSPS); a grant from the Core Center for iPS Cell Research (JP21bm0104001), Research Center Network for Realization of Regenerative Medicine from Japan Agency for Medical Research and Development (AMED); a grant from the Takeda Science Foundation; and the iPS Cell Research Fund from the Center for iPS Cell Research and Application, Kyoto University.

## Author contributions

Conceptualization, K.T., K.W.; Methodology, K.T., K.W.; Investigation, K.T., C.O., M.N., M.I., K.Y., Y.M.; Formal analysis, K.T., C.O., M.I., T.T.; Writing, K.T., C.O., M.I., K.W.; Supervision, K.T., S.Y.

---

**Fig. S1.**
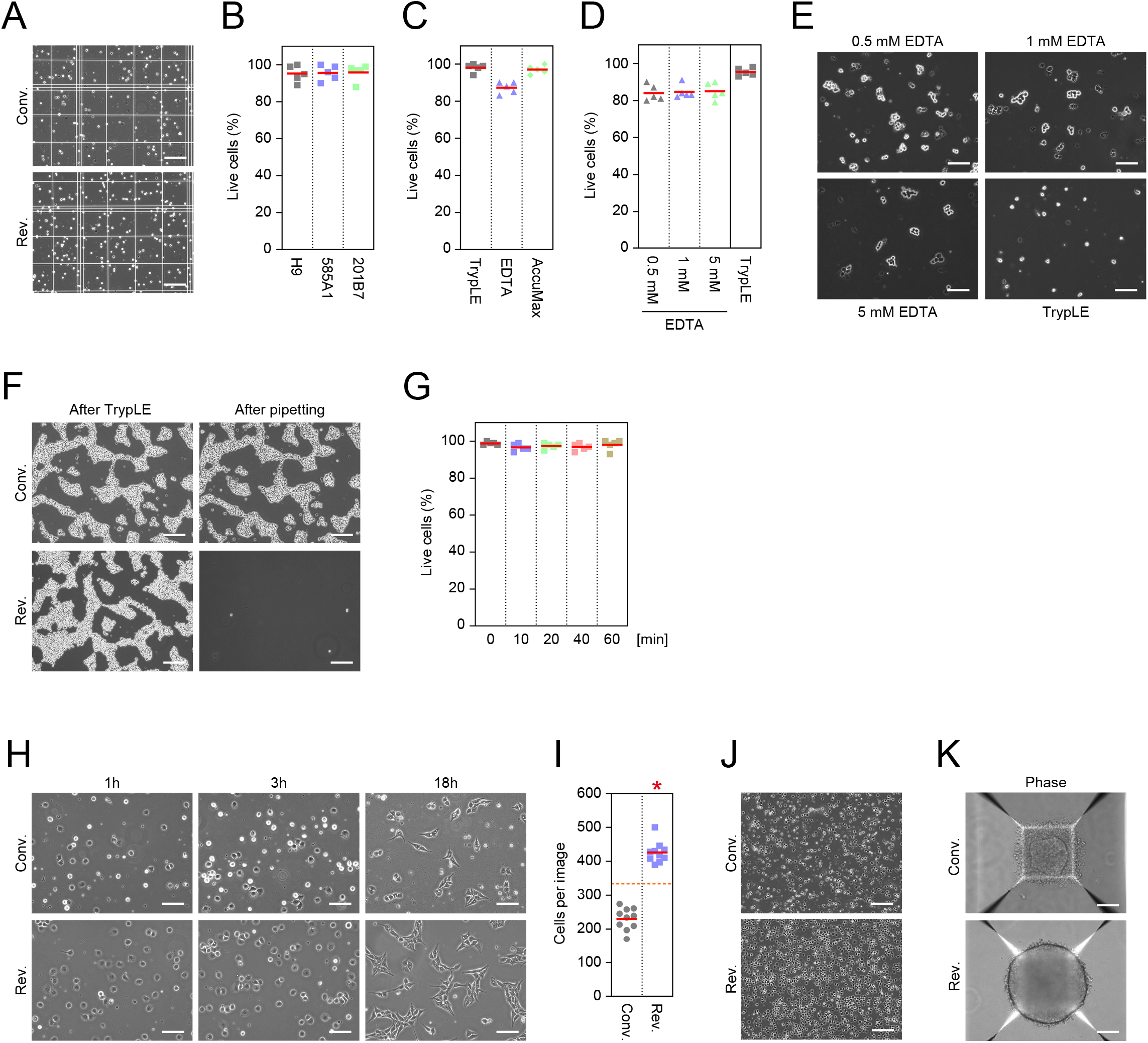
Supportive data of the revised passaging method, related to Fig. 1. A. Representative images of trypan blue-stained cells on a hematocytometer. Bars, 200 μm. B. The viability of three independent PSC lines after harvesting by the revised method. n=5. C. The viability of PSCs maintained on vitronectin in E8 media after harvesting. n=5. D. The viability of PSCs after harvesting with EDTA (0.5-5 mM) or TrypLE. n=5. E. Representative images of EDTA-treated cells after dissociation. Bars, 100 μm. F. Representative images of PSCs before and after dissociation by pipetting in growth media (Conv.) and TrypLE (Rev.). Shown are cells just after TrypLE treatment (left) and after subsequent dissociation by 10 times pipetting and washing the floating cells out (right). Bars, 200 μm. G. The viability of PSCs after incubation at the indicated times in TrypLE. n=5. H. Representative images of PSCs at 2 × 10^5^ cells per well of a 6-well plate after the indicated times of plating. Bars, 100 μm. I. The cell number after 18 h of plating. Microscopic views under a 10x objective were randomly imaged, and the number of nuclei labeled by Hoechst 33342 were counted. *p<0.05 vs. Conv., unpaired t-test. n=10. J. Representative images of PSCs at 1 million cells per well of a 6-well plate after 1 h of plating. Bars, 200 μm. K. Representative images of neurospheres after 18 h of plating in AggreWell800. These are the phase contrast images of the fluorescent images shown in Fig. 1M. Bars, 100 μm.

**Fig. S2.**
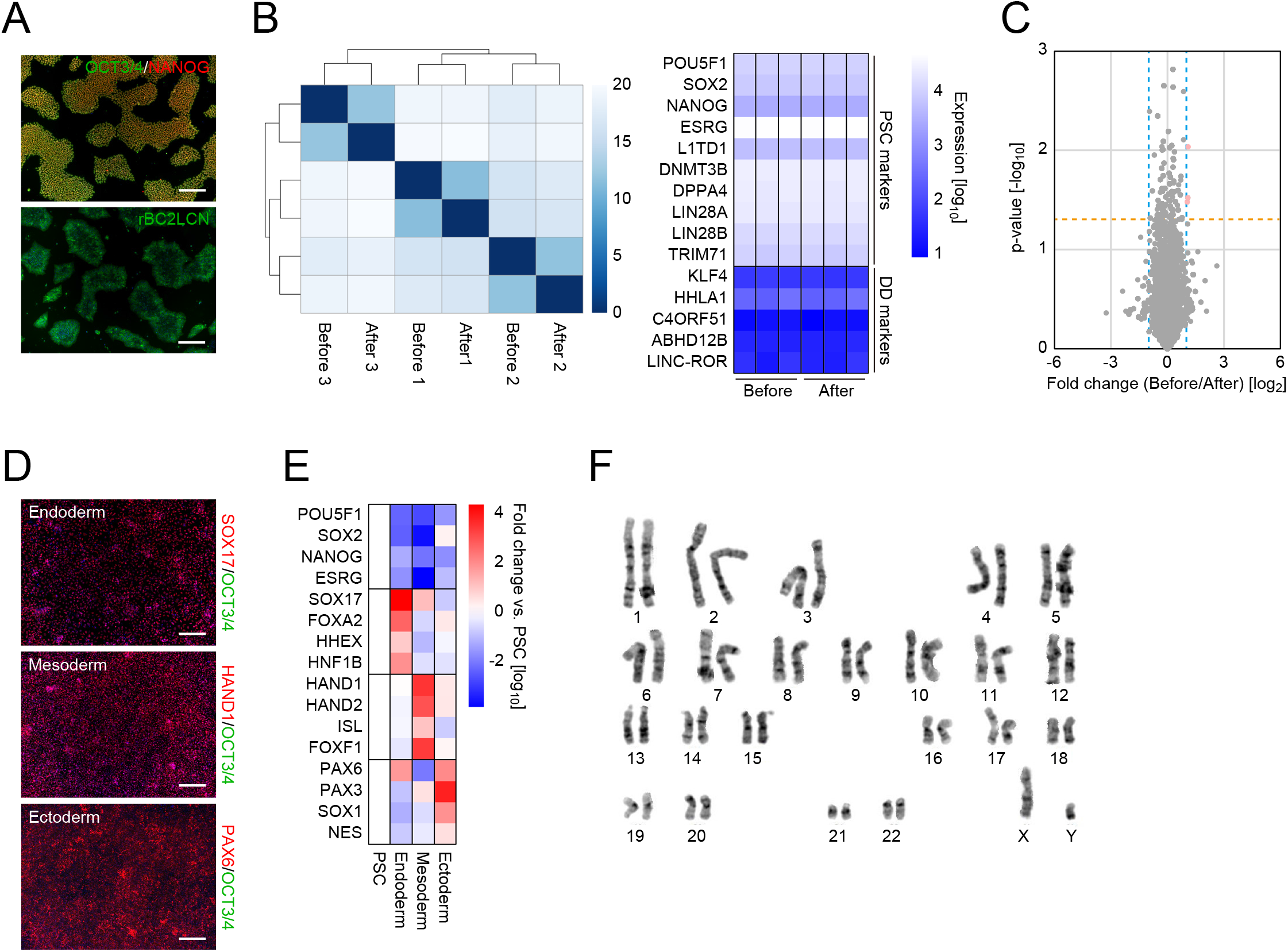
WTC11 iPSCs after 30 passages with the revised method, related to Fig. 2. A. PSC markers in WTB6 iPSCs after 30 passages with the revised method. Nuclei were visualized with Hoechst 33342 (blue). Bars, 200 μm. B. The RNA expression. Shown are clustering analysis of global RNA expression (left) and the PSC and differentiation-defective (DD) markers between WTC11 iPSCs before and after ≥30 passages with the revised method. n=3. C. The protein expression. The volcano plot shows the comparison of global protein expression between WTC11 iPSCs before and after cultivation with passaging using the revised method (>30 passages). n=3. The mandarin-colored and turquoise-colored broken lines indicate p=0.05 and fold change=2, respectively. Red dots indicate differentially expressed proteins such as MREG, PLCD3 and STK19. D. Directed differentiation into trilineage. Shown are representative immunocytochemistry images of differentiated cells stained for each lineage marker, such as SOX17, HAND1 and PAX6 (red), and a PSC marker, OCT3/4 (green). Nuclei were visualized with Hoechst 33342 (blue). Bars, 200 μm. E. The expression of lineage markers. The heatmap shows the relative expression of PSC and lineage markers in directed differentiated cells compared to parental iPSCs. n=3. F. A representative G-banding image showing normal 46XY karyotype.

**Table S1.**
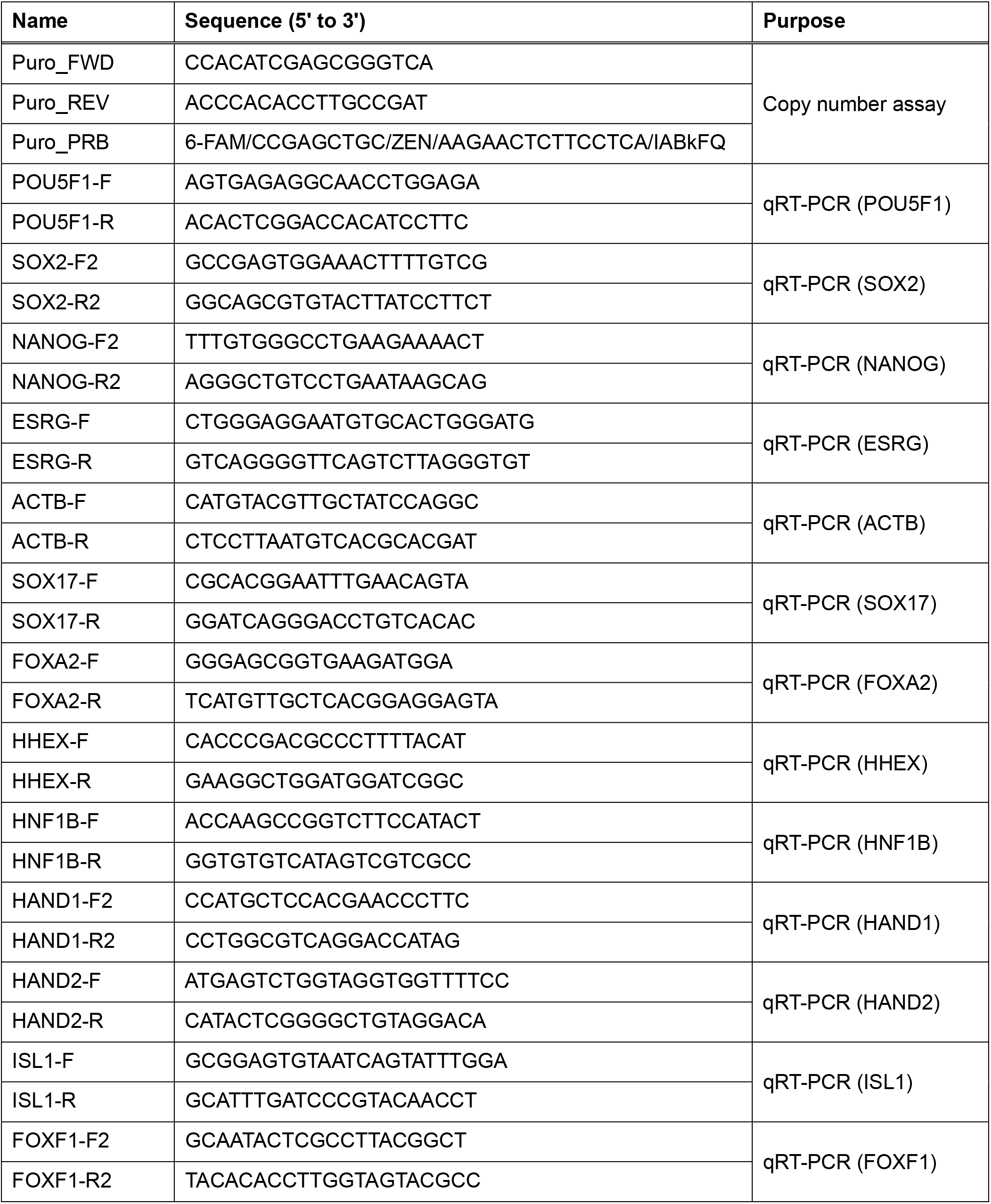
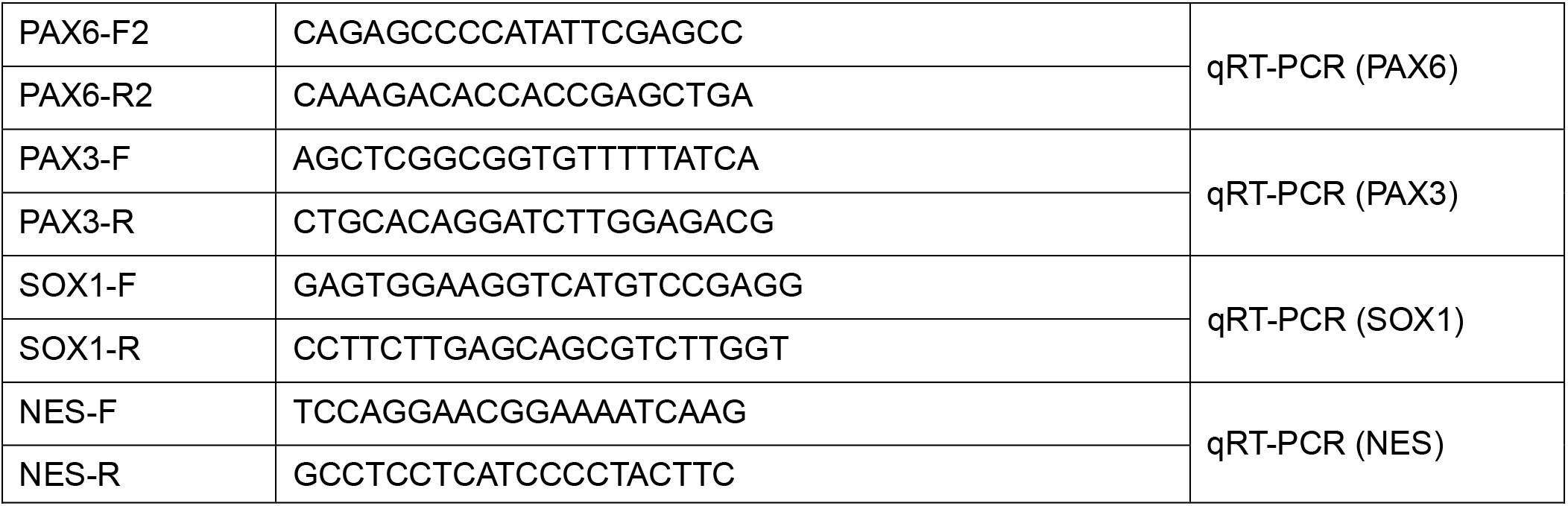
The sequences of the oligonucleotides used in this study.

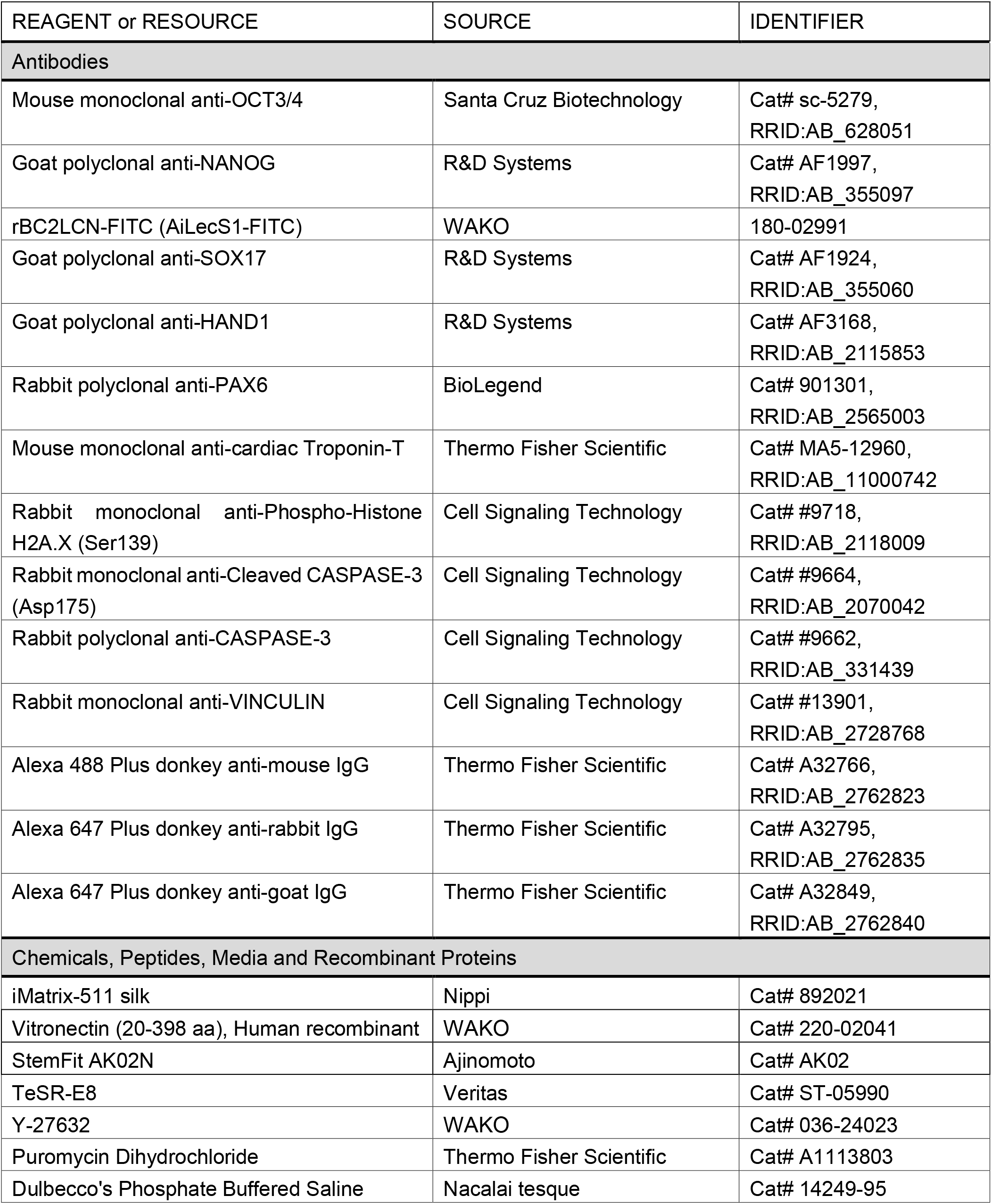
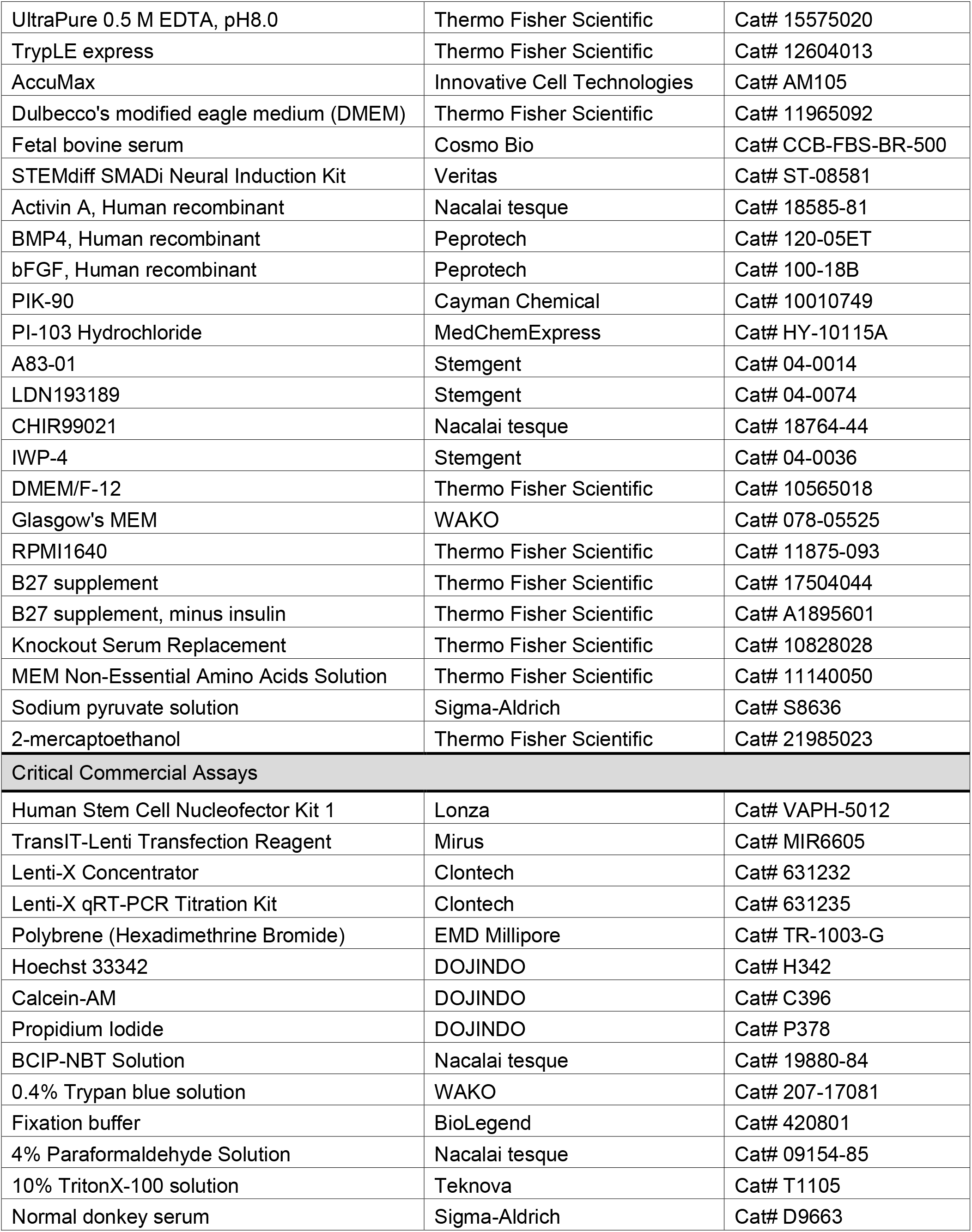
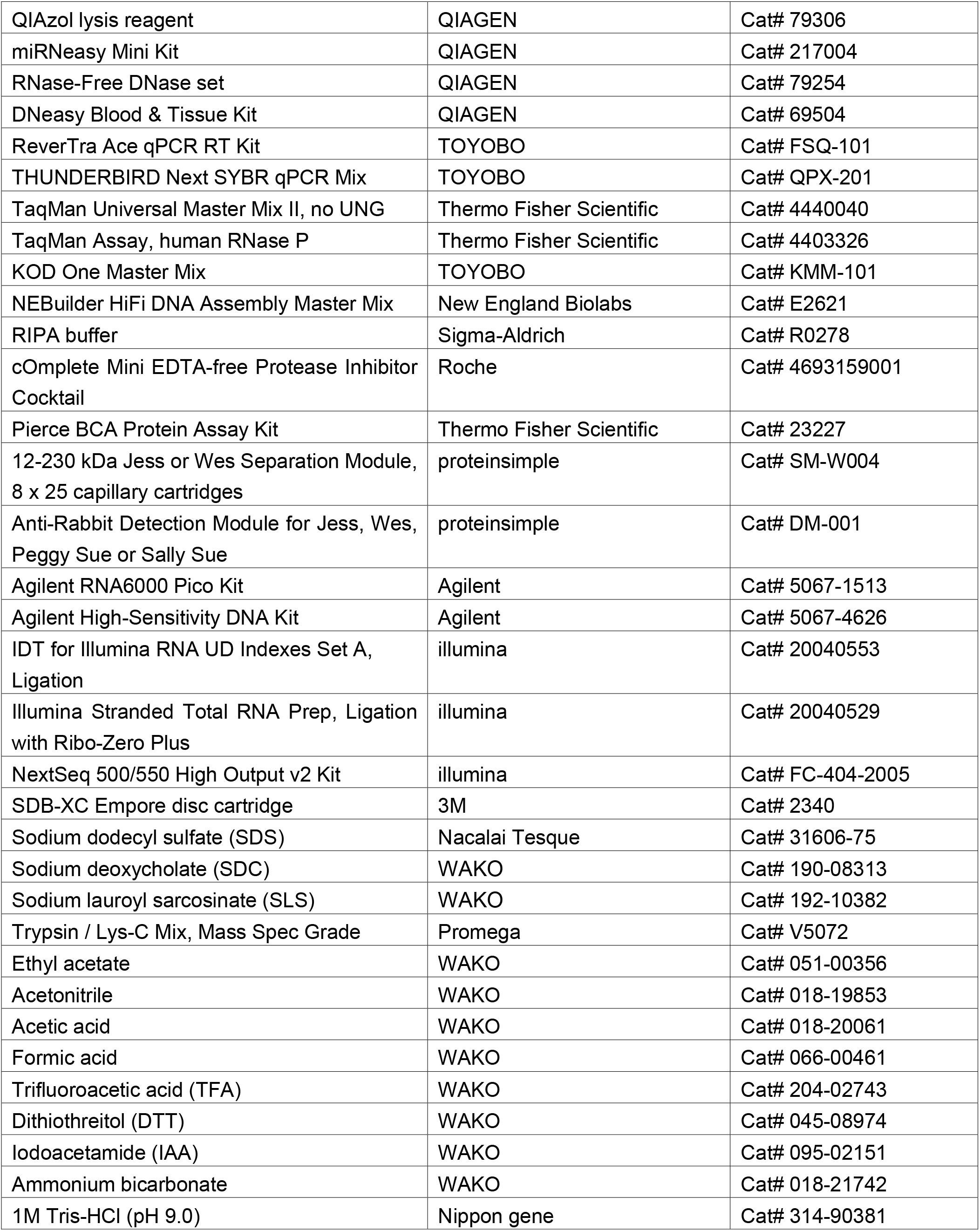
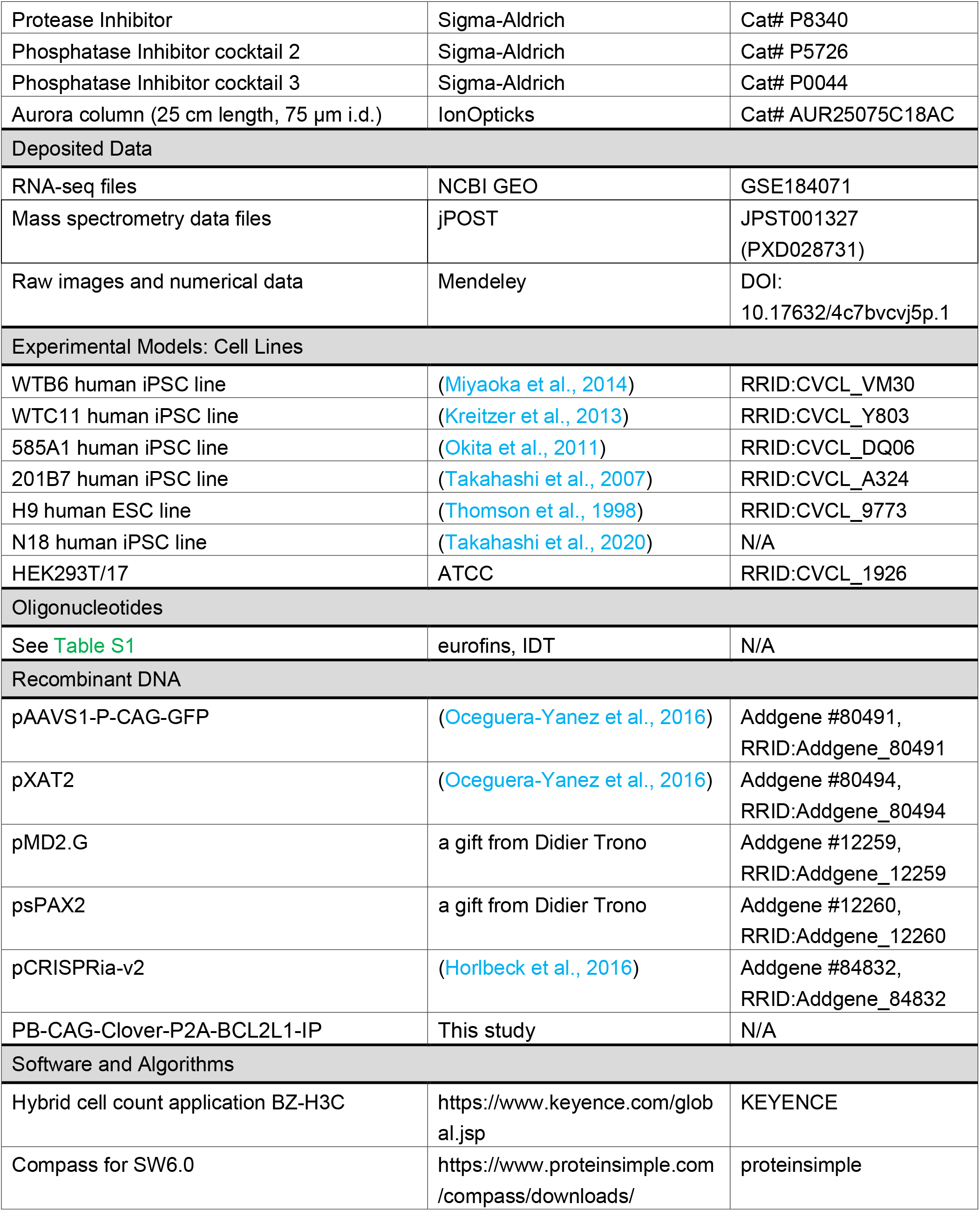
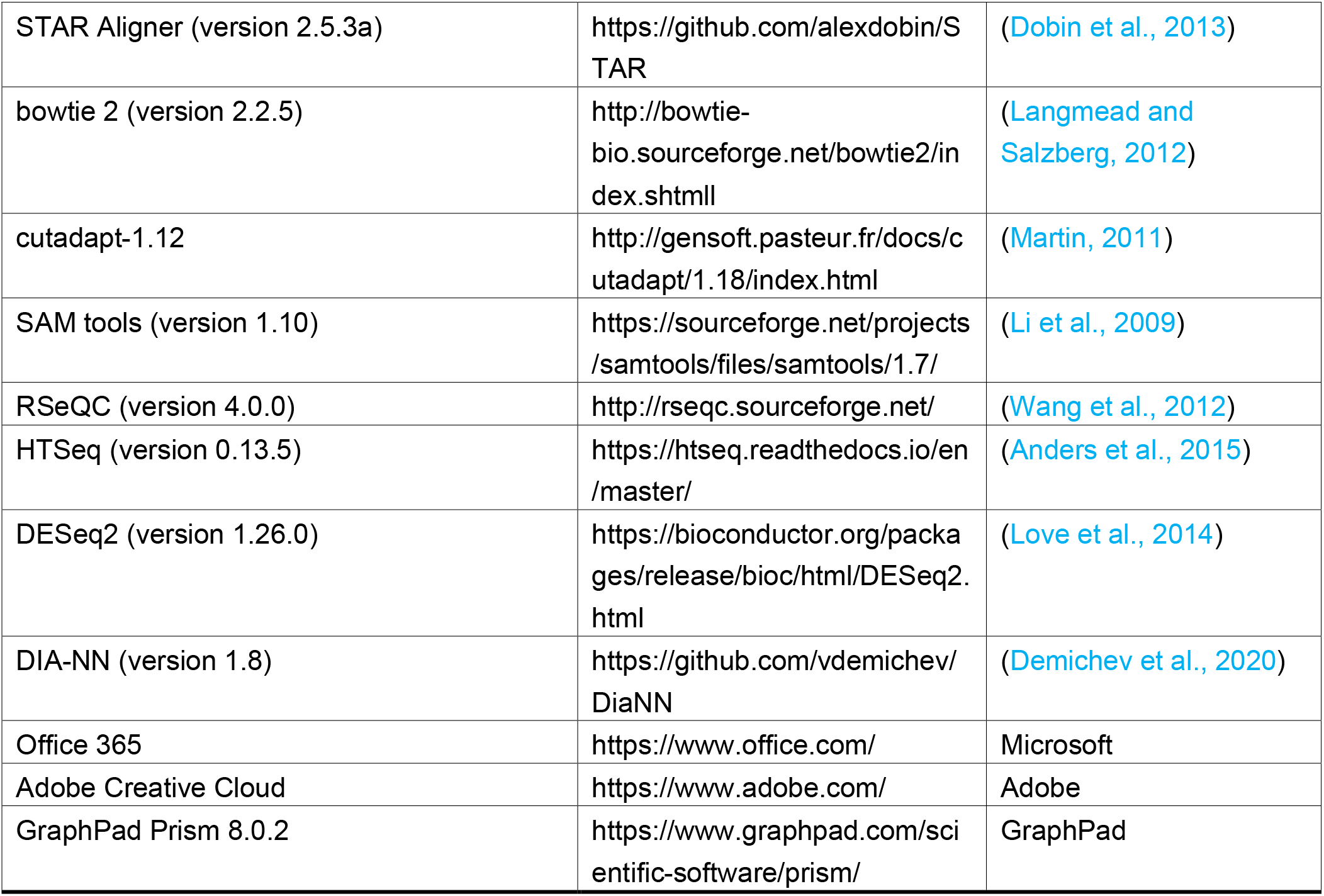
KEY RESOURCES TABLE

## RESOURCE AVAILABILITY

### Lead Contact

Further information and requests should be directed to the lead contact, Kazutoshi Takahashi

### Materials Availability

Unique reagents generated in this study are available from the lead contact with a Materials Transfer Agreement.

### Data and Code Availability

RNA sequencing results and proteome data are accessible in the Gene Expression Omnibus database of the National Center for Biotechnology Information website (accession number: GSE184071) and in the Japan Proteome Standard Repository/Database (accession number: JPST001327), respectively. The images and numerical data and statical analysis results that were not shown in the paper are available at Mendeley (DOI: 10.17632/4c7bvcvj5p.1).

### Experimental model and subject details

The human pluripotent stem cell (PSC) lines WTB6 iPSC (P49) (RRID:CVCL_VM30), WTC11 iPSC (P46) (RRID:CVCL_Y803), 585A1 iPSC (P49) (RRID:CVCL_DQ06), 201B7 iPSC (P28) (RRID:CVCL_A324), and H9 ESC (P51) (RRID:CVCL_9773) along with HEK293T/17 (P26) (RRID:CVCL_1926) were cultured in humidified incubators at 37°C in 5% CO_2_ and 20% O_2_. All reagents were warmed in a water bath set at 23°C before use unless otherwise noted. We confirmed all cell lines used in the study were negative for mycoplasma contamination by performing a periodical test (Young et al., 2010). The karyotype analysis was performed by Nihon Gene Research Laboratories, Inc. The experiments using H9 ESCs were conducted in conformity with “The Guidelines on the Distribution and Utilization of Human Embryonic Stem Cells” of the Ministry of Education, Culture, Sports, Science and Technology, Japan. WTB6 and WTC11 were provided by Bruce R. Conklin (The Gladstone Institutes and University of California San Francisco).

### Method details

#### Cell culture

Human PSCs listed in the Key Resource Table were maintained on a laminin 511 E8 fragment (iMatrix, Nippi) in StemFit AK02N media (Ajinomoto) as described previously (Miyazaki et al., 2012; Nakagawa et al., 2014). We also cultured human PSCs on recombinant human vitronectin (20-388 a.a.) (rhVTN, WAKO) in TeSR-E8 media (Veritas) (Beers et al., 2012; Chen et al., 2011). The media was changed daily. HEK293T/17 cells (ATCC) were grown in Dulbecco’s modified eagle medium (DMEM, Thermo Fisher Scientific) containing 10% fetal bovine serum (FBS, Cosmo Bio).

#### The revised passaging method

In the study, after three days of seeding human PSCs at 2 × 10^5^ cells per well of a 6-well plate (~20,000 cells/cm^2^), we performed the passaging with the indicated method and reagent unless otherwise stated. The cells were washed once with Dulbecco’s Phosphate-Buffered Saline, calcium-free and magnesium-free (hereafter D-PBS, Nacalai tesque) and treated with a dissociating solution, such as TrypLE express (Thermo Fisher Scientific), 0.5-5 mM EDTA, which was prepared by diluting UltraPure 0.5 M EDTA (Thermo Fisher Scientific) in D-PBS, or AccuMax (Innovative Cell Technologies), for 10 min at 37°C. Then the dissociation was performed in the detachment reagent by pipetting up and down using a 1-mL micropipette or using a cell scraper (IWAKI), and the cell suspension was mixed with StemFit AK02N (~10 times volume of dissociation reagent) in a 15-mL tube. Live and dead cells were counted by mixing 1:1 with 0.4% trypan blue solution (WAKO) and using a TC20 automated cell counter (Bio-Rad). After counting the cell number, the cells were spun down at 240 xg for 5 min at room temperature, and the cell pellet was resuspended in StemFit AK02N media. A Rho associate kinase inhibitor, Y-27632 (10 μM, WAKO), and iMatrix (0.25 μg per 1 cm^2^ of the culture vessel surface area) were added to the media during the replating (Miyazaki et al., 2017). For the culture system of TeSR-E8 media and rhVTN, the culture vessels were coated with rhVTN (0.75 μg per 1 cm^2^) for 1 h at room temperature before use. We routinely plated 2 × 10^5^ live cells per well of a 6-well plate and passaged them every three days.

#### Conventional passaging

The conventional passaging method was described previously (Nakagawa et al., 2014). In brief, the cells were washed with D-PBS and treated with a 1:1 mix of TrypLE express and 0.5 mM EDTA (T1E1) for 5 min at 37°C. Then we aspirated off the supernatant, added an appropriate volume of StemFit AK02N media, harvested the cells using a cell scraper, and collected them in a 15-mL tube. We dislodged the cells using a cell scraper within 30 s after adding the media unless otherwise noted. After that, we counted the cells and plated 2 × 10^5^ live cells per well of a 6-well plate in StemFit AK02N media supplemented with 10 μM Y-27632 and iMatrix (0.25 μg per 1 cm^2^ of the culture vessel surface area). To fairly compare the two methods, we incubated the cells for 10 min rather than 5 min in TrypLE instead of T1E1 in the experiments shown in the figures except Figs. 1B and 1C.

#### Cell adhesion test

Cells harvested using the indicated dissociation protocol were plated at 2 × 10^5^ cells per well of a 6-well plate by the non-coating method. Eighteen hours after plating, the cells were washed three times with PBS and fixed by treating them with 4% paraformaldehyde (BioLegend) for 15 min at room temperature. Then the fixation buffer was replaced with D-PBS supplemented with 1 μg/mL Hoechst 33342 (DOJINDO) and incubated for 60 min at room temperature with protection from light. Ten images of each sample were randomly taken using a BZ-X710 all-in-one fluorescence microscope (KEYENCE), and the number of Hoechst-stained nuclei was counted using a hybrid cell count application BZ-H3C (KEYENCE).

#### Single cell plating in clonal density

The cells were harvested using the methods indicated above and serially diluted to prepare the cell suspension at 50 live cells/mL in StemFit AK02N plus 10 μM Y-27632 and iMatrix (0.25 μg per 1 cm^2^ of the culture vessel surface area). Nine milliliters of the cell suspension containing 450 cells were transferred onto a 100-mm dish and incubated overnight at 37°C. The media was changed every other day with fresh StemFit AK02N. After 8 days of culture, the cells were fixed with 4% paraformaldehyde solution (Nacalai tesque) for 2 min at room temperature and incubated with BCIP-NBT solution (Nacalai tesque) for 60 min to visualize alkaline phosphatase positive (+) colonies.

#### Live and dead cell staining

The cells were treated with 1 μg/mL of Calcein-AM (DOJINDO) and 2 μg/mL of propidium iodide (PI, DOJINDO) for 15 min at 37°C to discriminate between live and dead cells. After the incubation, the cells were imaged using the BZ-X710 with a 10x objective, and the number of PI (+) cells was counted using the BZ-H3C.

#### RNA isolation and quantitative reverse transcription polymerase chain reaction

The cells were lysed with a QIAzol reagent (QIAGEN), and the total RNA was extracted using a miRNeasy Mini Kit (QIAGEN) with on-column DNase treatment (QIAGEN) according to the manufacturer’s protocol. One microgram of purified RNA was applied for the first-strand cDNA synthesis using a ReverTra Ace qPCR RT Master Mix (TOYOBO). Quantitative reverse transcription polymerase chain reaction (qRT-PCR) was performed using a THUNDERBIRD Next SYBR qPCR Mix (TOYOBO) and gene specific primers on a QuantStudio 3 real-time PCR system (Applied Biosystems). The raw Ct values were normalized to the housekeeping gene human ACTB via the double delta Ct method, and then the relative expression was calculated as the fold-change from the control. Primer sequences are listed in Table S1.

#### Western blotting

Cells harvested using the conventional or revised method were pelletized by centrifugation and washed once with D-PBS. The pellets were lysed with RIPA buffer (Sigma-Aldrich) supplemented with complete EDTA-free protease inhibitor cocktail (Sigma-Aldrich) and stored at −80°C until use. The cell lysates were heated at 95°C for 5 min, spun at 15,300 xg for 15 min at 4°C, and the cleared supernatant was collected and used for the following analysis. The protein concentration was measured using a Pierce BCA Protein Assay Kit (Thermo Fisher Scientific) and EnVision 2104 plate reader (Perkin Elmer) as instructed. We utilized a size-based protein analysis on a Wes automated capillary electrophoresis platform (proteinsimple) and 12-230 kDa Separation Module (proteinsimple) as instructed. We loaded 2.0-2.4 μg of cell lysates for each detection along with the following antibodies: rabbit monoclonal anti-phospho-H2AX (Ser139) (1:50, Cell Signaling Technology), rabbit polyclonal anti-CASPASE-3 (1:50, Cell Signaling Technology), rabbit monoclonal anti-cleaved CASPASE-3 (1:50, Cell Signaling Technology), and rabbit monoclonal anti-VINCULIN (1:250, Cell Signaling Technology). The data was visualized and analyzed using Compass for SW6.0 software (proteinsimple).

#### Lentiviral production and transduction

One day before the transfection, we plated HEK293T/17 cells at 5 × 10^6^ cells per collagen I-coated 100-mm dish (IWAKI) in DMEM containing 10% FBS. The next day, we transfected 5 μg of pCRISPRia-v2 (a gift from J. Weissman), 3.75 μg of psPAX2 (a gift from D. Trono) and 1.25 μg of pMD2.G (a gift from D. Trono) using TransIT-Lenti Transfection Reagent (Mirus) according to the manufacturer’s instruction. Forty-eight hours after the transfection, we collected the virus-containing supernatant and filtered it through a 0.45-μm pore size PVDF filter (EMD Millipore) to remove the cell debris. The filtered supernatant was mixed 3:1 with Lenti-X Concentrator (Clontech) and incubated at 4°C overnight. The mixture was centrifuged at 1,500 xg for 45 min at 4°C. After aspirating the supernatant off, the tube was centrifuged at 1,500 xg for 5 min at 4°C, and the rest of the supernatant was completely removed using a micropipette. Then the pellet was resuspended in D-PBS (1/100 of the initial volume of virus-containing supernatant) and stored in aliquots at −80°C until use. The number of viral particles (vps) was quantified using a Lenti-X qRT-PCR Titration Kit (Clontech) according to the manufacturer’s protocol. The cells plated at 1 × 10^6^ cells per well of a 6-well plate one day before infection were incubated with 1,000 vps/cell (multiplicity of infection = 1) along with 8 μg/mL polybrene at 37°C for 18 h. The next day, the virus-containing media was replaced with fresh StemFit AK02N media. On day 3 post-infection, the cells were selected with 0.5 μg/mL puromycin (Thermo Fisher Scientific) until non-transduced cells died completely.

#### Targeting

The CRISPR/Cas9-mediated knock-in experiment was performed as described previously (Oceguera-Yanez et al., 2016; Takahashi et al., 2020) with slight modifications. In brief, we transfected 3 μg of pXAT2 and 7 μg of pAAVS1-P-CAG-GFP into 1 million WTB6 iPSCs using a Human Stem Cell Nucleofector Kit I (Lonza) and the A-023 program of a Nucleofector IIb device (Lonza). Fifty thousand electroporated cells were plated onto a 100-mm dish in StemFit AK02N supplemented with 10 μM Y-27632 and iMatrix (0.25 μg per 1 cm^2^ of the culture vessel surface area). Three days after the electroporation, we started the selection with 0.5 μg/mL puromycin and continued it until the non-transfected control died completely and the colonies of the targeted cells grew enough. The colonies were visualized by staining using BCIP-NBT solution, and the number of alkaline phosphatase (+) colonies were counted. In this assay, we considered drug-resistant colonies are AAVS1-targeted clones.

#### Plasmid construction

A triple tandem repeat of Simian Virus 40 nuclear localization signal-tagged Clover (with no stop codon) and a protein coding sequence of human BCL2L1 gene linked in-frame by the porcine teschovirus-derived 2A peptide sequence were generated by PCR using KOD One Master Mix (TOYOBO). The fragment was inserted into the XhoI site of PB-CAG-IP using NEBuilder HiFi DNA Assembly (New England Biolabs) and verified by sequencing. The resulting plasmid encodes Clover, BCL2L1 and puromycin-resistance gene driven by a constitutively active CAG promoter. The sequence can be provided upon request.

#### Generation of BCL2L1-expressing iPSC line

We transfected 7 μg of PB-CAG-Clover-P2A-BCL2L1-IP and 3 μg of pCW-hyPBase into one million WTB6 iPSCs as described previously (Takahashi et al., 2020; Yusa et al., 2011). Two days after the transfection, the cells were selected with 0.5 μg/mL of puromycin until the non-transfected cells died completely. The colonies expressing Clover uniformly were chosen and isolated.

#### Cell competition assay

WTB iPSCs expressing exogenous BCL2L1 were mixed at a 1:19 ratio with WTB6 parental iPSCs. The mixed cells were plated at 2 × 10^5^ cells per well of a 6-well plate and passaged every three days as described above. The cell population was analyzed by the microscopic observation of Clover fluorescence and PCR-based copy number assay for a puromycin-resistance gene.

#### Genomic PCR for copy number quantification

Genomic DNA was isolated using a DNeasy Blood & Tissue Kit (QIAGEN) according to the manufacturer’s protocol. Thirty nanograms of purified DNA was used for qPCR using TaqMan Universal Master Mix II, no UNG (Thermo Fisher Scientific) on a QuantStudio 3 Real Time PCR System. A PrimeTime qPCR Assay (Integrated DNA Technologies) was used to detect the puromycin-resistance gene in the pCRISPRia-v2 lentiviral vector or PB-CAG-IP vector, and a TaqMan Copy Number Reference Assay human RNase P (Thermo Fisher Scientific) was used as an internal control. For quantification of the lentiviral insertion, we used the genomic DNA of the N18 iPSC line carrying one copy of the puromycin-resistance gene as a control (Takahashi et al., 2020). The primer and probe sequences are provided in Table S1.

#### Endoderm differentiation

The directed differentiation into endoderm was performed as described previously with slight modifications (Loh et al., 2014; Martin et al., 2020). A day before the differentiation, human PSCs were plated at 5 × 10^5^ cells per well of a 12-well plate on iMatrix in StemFit AK02N plus 10 μM Y-27632. The cells were treated with 100 ng/mL Activin A (Nacalai tesque), 3 μM CHIR99021 (Nacalai tesque), 20 ng/mL bFGF (Peprotech) and 50 nM PI-103 (Cayman Chemical) in differentiation media 1 (DM1), which consisted of DMEM/F12 (Thermo Fisher Scientific), 2% B27 supplement (Thermo Fisher Scientific), 1% MEM Non-Essential Amino Acids (NEAA, Thermo Fisher Scientific) and 0.1 mM 2-mercaptoethanol (Thermo Fisher Scientific) for 24 h (day 1). Then the cells were cultured in DM1 supplemented with 100 ng/mL Activin A and 250 nM LDN193189 for 48 h (days 2 and 3). Two days later, the cells were maintained in DM1 containing 100 ng/mL Activin A (days 4 and 5). The media was changed daily.

#### Mesoderm differentiation

Mesoderm differentiation was performed as described previously with slight modifications (Loh et al., 2016; Martin et al., 2020). A day before the differentiation, human PSCs were plated at 5 × 10^5^ cells per well of a 12-well plate on iMatrix in StemFit AK02N plus 10 μM Y-27632. The cells were cultured in DM1 supplemented with 30 ng/mL Activin A, 40 ng/mL BMP4 (Peprotech), 6 μM CHIR99021, 20 ng/mL bFGF and 100 nM PIK-90 (MedChemExpress) for 24 h (day 1). Then the cells were cultured in DM1 supplemented with 40 ng/mL BMP4, 1 μM A83-01 and 4 μM CHIR99021 for 48 h (days 2 and 3). Two days later, the cells were maintained in DM1 containing 40 ng/mL BMP4 (days 4 and 5). The media was changed daily.

#### Ectoderm Differentiation

The directed differentiation into neuroectoderm was performed as described previously (Chambers et al., 2009; Doi et al., 2014; Takahashi et al., 2020). A day before the differentiation, human PSCs were plated at 5 × 10^5^ cells per well of a 12-well plate on iMatrix in StemFit AK02N plus 10 μM Y-27632. The cells were treated with 1 μM A83-01 and 250 nM LDN193189 in Glasgow’s MEM (WAKO) containing 8% Knockout serum replacement (Thermo Fisher Scientific), 1 mM sodium pyruvate (Sigma-Aldrich), 1% NEAA and 0.1 mM 2-mercaptoethanol for 5 days. The media was changed daily.

#### Cardiomyocyte differentiation

We performed directed cardiomyocyte differentiation as described previously (Lian et al., 2012) with slight modification. Two days before starting the differentiation, cells harvested using the conventional or revised method were plated at 3.5 × 10^5^ cells per well of a 12-well plate in StemFit AK02N supplemented with 10 μM Y-27632 and iMatrix (0.25 μg per 1 cm^2^ of the culture vessel surface area). On the day we designated as day 0, the media was replaced with RPMI1640 supplemented with 2% B27 supplement (minus insulin) and 4 μM CHIR99021. On day 1, we removed CHIR99021. On day 3, we added 5 μM IWP-4 (Stemgent) until day 5. On day 7, we switched the media to RPMI containing B27 supplement (plus insulin) and cultured the cells until day 12.

#### Neurosphere formation

Cells harvested using the conventional or revised method were transferred at 3 × 10^6^ cells per well of an AggreWell 800 24-well plate (Veritas) in STEMdiff Neural Induction Medium supplemented with SMAD inhibitor (Veritas) and 10 μM Y-27632. Then the plate was briefly spun at 100 xg for 3 min and incubated at 37°C for 18 h. After the incubation, we performed Calcein/PI staining as described above. To measure the size of the aggregates, we imaged Calcein (+) cells under a 488 nm wavelength filter and measured the diameter of each aggregate using the BZ-H3C. We only evaluated single spheres in a microwell of AggreWell 800 plate.

#### Immunocytochemistry

Indirect immunocytochemistry was performed as described previously (Takahashi et al., 2021). The cells were washed once with D-PBS and fixed using 4% paraformaldehyde for 15 min at room temperature. The fixed cells were permeabilized and blocked with D-PBS containing 0.1% TritonX-100 (Teknova), 1% bovine serum albumin (BSA, Thermo Fisher Scientific) and 2% normal donkey serum (Sigma-Aldrich) for 45 min at room temperature. We incubated the samples with the primary antibody diluted in D-PBS containing 1% BSA at 4°C for 6-18 h. After washing with D-PBS three times, the samples were incubated with the secondary antibody diluted in D-PBS containing 1% BSA and 1 μg/mL Hoechst 33342 for 45-60 min at room temperature in the dark. After a brief wash with D-PBS, the cells were observed using a BZ-X710. The antibodies used in the study are as follows: mouse monoclonal anti-OCT3/4 (1 μg/mL, SantaCruz), goat polyclonal anti-NANOG (5 μg/mL, R&D systems), goat polyclonal anti-SOX17 (0.5 μg/mL, R&D systems), goat polyclonal anti-HAND1 (5 μg/mL, R&D systems), rabbit polyclonal anti-PAX6 (2 μg/mL, BioLegend), mouse monoclonal anti-cardiac Troponin-T (5 μg/mL, Thermo Fisher Scientific), Alexa 488 Plus donkey anti-mouse IgG (4 μg/mL, Thermo Fisher Scientific), Alexa 647 Plus donkey anti-rabbit IgG (4 μg/mL, Thermo Fisher Scientific) and Alexa 647 Plus donkey anti-goat IgG (4 μg/mL, Thermo Fisher Scientific).

#### rBC2LCN staining

The cells were washed once with D-PBS and fixed using 4% paraformaldehyde for 15 min at room temperature. The fixed cells were incubated in D-PBS supplemented with 1/100 volume of rBC2LCN-FITC (WAKO) and 1 μg/mL Hoechst 33342 for 30 min at 37°C with protection from light. Then the cells were washed three times with D-PBS and observed using the BZ-X710.

#### RNA sequencing (RNA-seq) and data analysis

For the RNA-seq analysis, we used WTB6 at early passage (P49, 50 and 51) and late passage (P49+30, 49+31 and 49+32) and WTC11 at early passage (P45, 46 and 47) and late passage (P45+30, 45+31 and 45+32). Cells were lysed using QIAzol reagent, and total RNA was purified as described above. Purified RNA samples were evaluated using an Agilent RNA6000 Pico Kit (Agilent) on a Bioanalyzer 2100 (Agilent). We performed the library preparation and subsequent analysis as described previously (Okubo et al., 2021a; Okubo et al., 2021b). In brief, 100 ng of purified RNA was applied to the library construction using the Illumina Stranded Total RNA Prep, Ligation with Ribo-Zero Plus (Illumina). The libraries evaluated using an Agilent High-Sensitivity DNA Kit (Agilent) were sequenced using a NextSeq 500/550 High Output v2 Kit (Illumina). We trimmed the adapter sequence using cutadapt-1.12 (Martin, 2011), excluded reads mapped to ribosomal RNA using SAM tools (version 1.10) and bowtie 2 (version 2.2.5) (Langmead and Salzberg, 2012; Li et al., 2009), aligned the reads to the hg38 human genome using STAR Aligner (Version 2.5.3a) (Dobin et al., 2013; Wang et al., 2012), used RSeQC (version 4.0.0) for the quality check, counted the reads using HTSeq (version 0.13.5) and GENCODE annotation file (version 35) (Anders et al., 2015; Frankish et al., 2019), and normalized the counts using DESeq2 (version 1.26.0) in R (version 3.6.1) (Love et al., 2014). A Wald test was performed using the DESeq2 package.

#### Proteome analysis

Cells were lysed with ice-cold PTS lysis buffer consists of 100 mM Tris-HCl (pH9.0) (Nippon Gene), 12 mM sodium deoxycholate (WAKO) and 12 mM sodium lauroyl sarcosinate (WAKO) supplemented with 1% phosphatase inhibitors (Sigma-Aldrich) and 1% protease inhibitor (Sigma-Aldrich). The lysates were subjected to reduction, alkylation, Lys-C/trypsin digestion (enzyme ratio: 1/100) and desalting, as previously described (Iwasaki et al., 2019). Two-hundred and fifty nanograms of the peptides were loaded and separated on an Aurora column (25 cm length, 75 μm i.d., IonOpticks) using a nanoElute (Bruker) for subsequent analysis by timsTOF Pro system (Bruker). The mobile phases were composed of 0.1% formic acid (solution A) and 0.1% formic acid in acetonitrile (solution B). A flow rate of 400 nL/min of 2-17% solution B for 60 min, 17-25% solution B for 30 min, 25-37% solution B for 10 min, 37-80% solution B for 10 min, and 80% solution B for 10 min was used (120 min in total). The applied spray voltage was 1400 V, and the interface heater temperature was 180°C. To obtain MS and MS/MS spectra, the Parallel Accumulation Serial Fragmentation (PASEF) acquisition method with data-independent acquisition (DIA) mode was used (diaPASEF) (Meier et al., 2020). For diaPASEF settings, 1.7 s per one cycle with precursor ion scan and 16 times diaPASEF scans were conducted with the MS/MS isolation width of 25 m/z, precursor ion ranges of 400−1200 m/z, ion mobility ranges of 0.57−1.47 V s cm^2^. The obtained DIA data were searched by DIA-NN (v1.8) (Demichev et al., 2020) against selected human entries of UniProt/Swiss-Prot release 2020_03 with the carbamidomethylation of cysteine as the fixed modification and protein N-terminal acetylation and methionine oxidation as the variable modification. For the other DIA-NN parameters, Trypsin/P protease, one missed cleavage, peptide length range of 7-30, precursor m/z range of 300-1800, precursor charge range of 1-4, fragment ion m/z range of 200-1800, and 1% precursor FDR were used. The values “PG.Normalised” from the results were used as representative protein area values for the comparison.

#### Statistics

Results are shown as individual data (colored dots) and means (red bars). We performed statistical analyses including unpaired, two-tailed t-tests to calculate p-values for the difference between the sample and control and one-way analysis of variance (ANOVA) for multiple comparisons using GraphPad Prism 8.0.2 (Graphpad) and Excel (Microsoft). p-values less than 0.05 were considered significant and are indicated by the asterisks in the figures.

